# Mechanical characteristics of ultrafast zebrafish larval swimming muscles

**DOI:** 10.1101/2020.04.02.010298

**Authors:** A. F. Mead, G. G. Kennedy, B. M. Palmer, A. M. Ebert, D. M. Warshaw

**Affiliations:** Department of Molecular Physiology and Biophysics, University of Vermont, Burlington, VT 05405; Department of Biology, University of Vermont, Burlington, VT 05405; Instrumentation and Model Facility, University of Vermont, Burlington, VT 05405

## Abstract

Zebrafish (*Danio rerio*) swim within days of fertilization, powered by muscles of the axial myotomes. Forces generated by these muscles can be measured rapidly in whole, intact larval tails by adapting protocols developed for *ex vivo* muscle mechanics. But it is not known how well these measurements reflect the function of the underlying muscle fibers and sarcomeres. Here we consider the anatomy of the 5-day-old, wild-type larval tail, and implement technical modifications to measuring muscle physiology in intact tails. Specifically, we quantify fundamental relationships between force, length, and shortening velocity, and capture the extreme contractile speeds required to swim with tail-beat frequencies of 80-100 Hz. Therefore, we analyze 1000 frames/second movies to track the movement of structures, visible in the transparent tail, which correlate with sarcomere length. We also characterize the passive viscoelastic properties of the preparation to isolate forces contributed by non-muscle structures within the tail. Myotomal muscles generate more than 95% of their maximum isometric stress (76±3 mN/mm^2^) over the range of muscle lengths used *in vivo*. They have rapid twitch kinetics (full width at half-maximum stress: 11±1 msec) and a high twitch to tetanus ratio (0.91±0.05), indicating adaptations for fast excitation-contraction coupling. Although contractile stress is relatively low, myotomal muscles develop high net power (134±20 W/kg at 80 Hz) in cyclical work loop experiments designed to simulate the *in vivo* dynamics of muscle fibers during swimming. When shortening at a constant speed of 7±1 muscle lengths/second, muscles develop 86±2% of isometric stress, while peak instantaneous power during 100Hz work loops occurs at 18±2 muscle lengths/second. These approaches can improve the usefulness of zebrafish as a model system for muscle research by providing a rapid and sensitive functional readout for experimental interventions.

**Statement of significance:** The zebrafish (*Danio rerio*) may prove a uniquely efficient model system for characterizing vertebrate muscle physiology. Transparent, drug-permeable larva – each, in essence, a fully functional muscle – can be generated rapidly, inexpensively, and in large numbers. Critically, the zebrafish genome contains homologs of major muscle genes and is highly amenable to manipulation. To reach its potential, reliable (and preferably rapid) means are needed to observe the effects of experimental interventions on larval muscle function. In the present study we show how mechanical measurements made on whole, intact larval tails can provide a readout of fundamental muscle-mechanical properties. Additionally, we show that these muscles are among the fastest ever measured, and therefore worthy of study in their own right.

## Introduction

Larvae of the zebrafish, *Danio rerio*, begin to move within 18 hours of fertilization, and swim freely when 3 days old (1, 2). The rapid development and early functionality of the muscles of the tail that power these movements have made zebrafish an important model to study vertebrate muscle development and disease (1-7). Due to the high degree of genetic and structural conservation among vertebrate muscles, a number of human myopathies (e.g. Duchenne muscular dystrophy) have an analogous phenotype in the zebrafish, making their larvae a model system of choice, based on several unique attributes (3-5, 8, 9). Importantly, larvae are transparent, enabling *in vivo* visualization of the underlying musculature, and are permeable to drugs, making them amenable to high-throughput studies (10-12). Tools for genetic modification are well developed, and the early functionality of the locomotory system often allows study of mutations and transgenes that would be embryonic-lethal in other vertebrate model systems (13-15).

Larval tail muscle fibers lie nearly parallel to the long axis of the tail and constitute the majority of its mass (16). Therefore, investigators have mounted intact tails between a force transducer and a servo motor for controlling tail length and then electrically stimulated the tails to contract (8, 17-19). By this approach, relationships have been characterized between force, length, and shortening velocity in tails at multiple stages of larval development, and in response to experimental perturbations (8, 17-19). The approach gives investigators with questions about the molecular determinants of muscle function easy access to the benefits of the zebrafish model. However, certain fundamental muscle-mechanics parameters derived from the mechanical behavior of the intact, mounted tail do not correlate with the extreme contractile speeds and power apparently required for fast larval swimming (20, 21). Could these differences in part be due to the simplifying assumption that the intact larval tail mechanics directly reflect the mechanics of the underlying musculature? Specifically, do changes in tail length equate to changes in the length of the muscle fibers and more importantly their constituent sarcomeres, the smallest contractile unit of muscle? In the present study, we sought to improve the usefulness of the zebrafish, intact larval tail preparation, by defining how tail- and sarcomere-level mechanics are related. Therefore, we used high speed imaging to track the position of structural features visible in the transparent larval tail that were then correlated to the underlying muscle fiber and sarcomere length. By this technique, we corrected for internal shortening within the tail so as to maintain truly isometric muscle fiber lengths during stimulated force development. In addition, given the presence of large, non-muscle structures within the tail (e.g. notochord) that must contribute to the tail’s mechanical properties, it was imperative that the passive mechanical properties of these non-muscle structures be defined before interpreting the origin of the tail’s force generation following electrical stimulation.

Through our approaches, we have established a baseline set of muscle mechanical measurements relating tail forces to true muscle fiber and sarcomere length and shortening velocity in the intact 5-day-old zebrafish tail, and measured mechanical power under approximated *in vivo* contractile conditions, using a modified work loop approach. Our results suggest that the tail muscle fibers generate their maximum force at the *in vivo* resting length of the tail and at these lengths, twitch force, with a 25 msec time course, nearly equals that generated during a tetanic contraction. Interestingly, the larval tail muscle fibers generate maximum power while shortening at velocities approaching 20 lengths/sec, a speed not previously observed in a vertebrate skeletal muscle. With such dynamic contractility, the myotomal muscles have the capacity to power the rapid bursts of swimming speed required to evade predators in the wild.

## Materials and Methods

### Zebrafish husbandry

All animals in this study were housed and used according to protocols approved by the Institutional Animal Care and Use Committee at the University of Vermont. Wildtype (AB-strain) zebrafish were maintained at 28° C in an institutional facility and mated to produce embryos for experimentation. Larvae were maintained in E3 medium (in mM: 5 NaCl, 0.17 KCl, 0.33 CaCl_2_, 0.33 MgSO_4_, plus 10^−5^ % Methylene Blue) and incubated at 28° C for five days prior to use. Unused larvae were anesthetized with tricane (0.02%) and euthanized on ice.

### Histology

Five day old larvae were euthanized by tricane overdose (0.05%) and fixed by immersion (0.1 M PIPES, 2.5% glutaraldehyde, 1% paraformaldehyde) at room temperature for one hour, then stored at 4° C. Larvae were then embedded in plastic (22) prior to sectioning. To visualize myotomal muscle morphology, larvae were oriented to obtain longitudinal- or cross-sections of muscles, and semithin sections (∼1 µm) were cut with glass knives on a Reichert Ultracut microtome, mounted on glass slides, stained with toluidine blue to highlight structural details. Light micrographs were obtained using a Zeiss Axiocam 208 (Carl Zeiss Microscopy LLC) or an Aperio VERSA 8 Scanning System (Leica Biosystems Imaging, Inc.) for whole slide imaging. To visualize individual sarcomeres, larvae were oriented as above and ultrathin sections (∼80 nm) were cut with a diamond knife. These were retrieved onto 200 mesh nickel grids and contrasted with uranyl acetate (2% in 50% ethanol) and lead citrate. Electron micrographs were obtained with a JEM 1400 transmission electron microscope (JEOL USA, Inc.) operating at 80kV and a bottom mounted AMT digital camera and software (Advanced Microscopy Techniques, Corp.).

Cross-sectional area (CSA) measurements of whole tails, fast myotomal muscles, and slow myotomal muscles were made by tracing features in 40X light micrographs of toluidine blue stained images. Anatomical landmarks for fast and slow myotomal muscle are well defined (23), which allowed us to outline each muscle type in ImageJ software (24) and calculate CSA with the software’s built-in area function. CSA values were measured in sections made at three points along the length of the tail corresponding to the center and approximate boundaries of tail sections when mounted for mechanical studies (see Figure 1) (n = 3).

**Figure 1.**
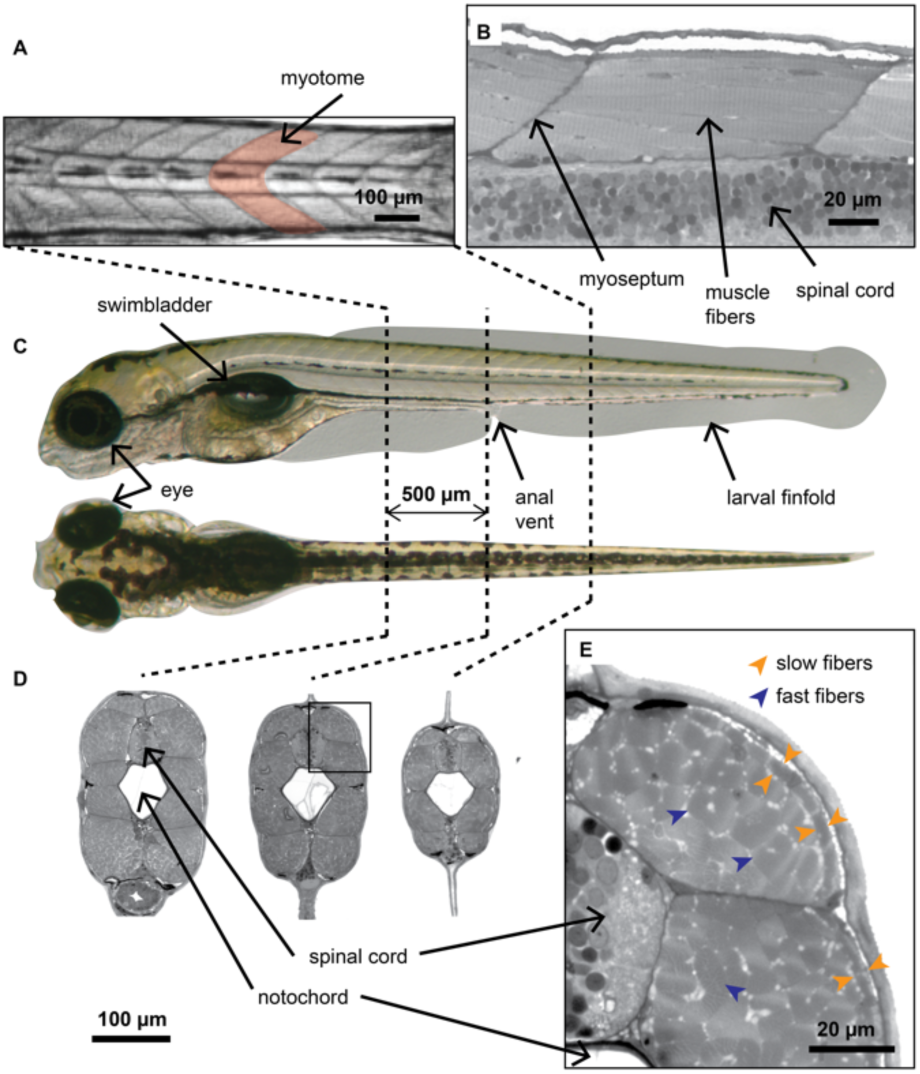
Anatomy of myotomal muscle in 5-day old zebrafish larvae. *(A)* A lateral bright field image of ∼1 mm long region of an intact tail used as a preparation in these studies (see Methods). The swimming muscles are arranged in a series of chevron-shaped myotomes *(shaded in pink). (B)* Longitudinal histological section of a single dorsal myotome stained to show structural detail. Muscle fibers span the myotome nearly parallel to the long axis of the tail and insert on planes of connective tissue *(myosepta)* that separate individual myotomes. A region of the spinal cord is also visible at the bottom of the image. *(C)* Dorsal and lateral views of a live larva showing major anatomical features. *(D)* Histological cross-sections from three points along the tail, ∼0.5 mm apart, showing the position of the notochord and spinal cord, which run the full length of the tail. *(E)* Detail from *D* (*black box*). Myotomal muscle makes up 67.3± 0.4 % (n=3) of tail CSA at this level. More than 90% of that area consists of fast fibers *(blue arrows)*. Slow muscle fibers at this age are confined to a single subcutaneous cell layer *(between orange arrows)*.

**Figure 2.**
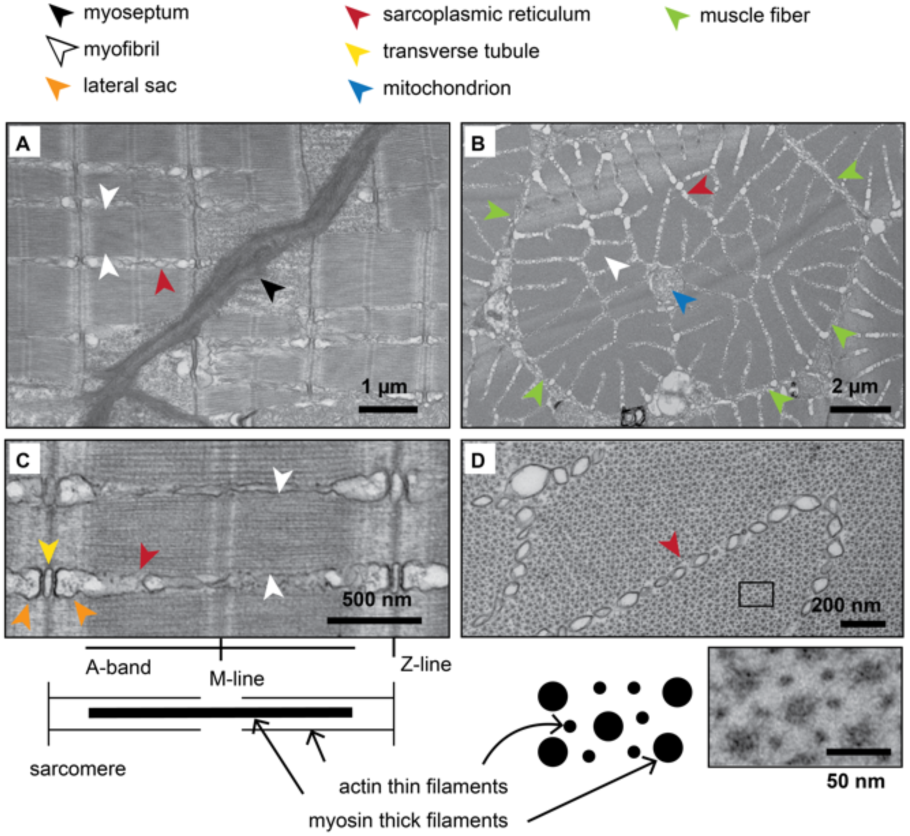
Fast muscle fiber ultrastructure. *(A)* Longitudinal TEM image of fast muscle. Individual myofibrils (*between white arrows*) bordered by sarcoplasmic reticulum *(red arrow)*, insert on the myoseptum *(black arrow)* at an angle consistent with the fiber orientation in Figure 1B. *(B)* Cross-sectional TEM image of fast muscle fibers. Single fiber (*border identified by green arrows*) has a centrally located mitochonrdrion *(blue arrow)*, and well distributed sarcoplasmic reticulum *(red arrow)* surrounding individual myofibrils ∼1 µm in diameter *(white arrow). (C)* Longitudinal TEM image at higher magnification. Sarcomeres (1.84±0.03 µm-long) within myofibrils (*between white arrows*) have clearly defined structural features: A-bands, M-lines, and Z-lines that relate to the positions of thin and thick filaments, as shown in an illustrated sarcomere below. Triad structures, consisting of a T-tubule *(yellow arrow)* and lateral sacs *(orange arrows)* are visible at the Z-lines, with sarcoplasmic reticulum *(red arrow)* bordering the myofibril. *(D)* Cross-sectional TEM image at higher magnification highlighting regular, hexagonal arrangement of thick (myosin) and thin (actin) filaments, as seen in inset below. Myofibrils are surrounded by sarcoplasmic reticulum (*red arrow*).

### Preparation of the larval tail for muscle mechanics experiments

Five day old larvae were selected at random, euthanized by tricane overdose (0.05% in E3), and transferred to Ringer’s solution (in mM: 117.2 NaCl, 4.7 KCl, 1.2 KH_2_PO_4,_ 1.2 MgCl_2_, 2.5 CaCl_2_, 25.2 NaHCO_3_, 11.1 glucose; oxygenated and equilibrated to pH 7.4 with a mixture of 95% O_2_ and 5% C0_2_)(17). Tails were removed by cutting with dissection scissors immediately caudal to the swim bladder (Fig. 3A), and mounted in a 2 ml Ringer’s-filled bath between a force transducer (KG4, SI Heidelberg) and linear servo motor/controller (MC1, SI Heidelberg) using spring clamps, as illustrated in Figure 3A and 3B. In order to study a consistent region of myotomal muscle, the clamps were set approximately 1 mm apart and tails were positioned such that the anal vent was equidistant from the attachment points (Figs. 1, 3B). The whole preparation was then slid into an experimental chamber within the bath, consisting of an optically clear quartz cylinder with an interior diameter of 3 mm and length of 10 mm, which enabled tighter temperature control and improved visualization. Mounted tails were illuminated in bright-field by a xenon arc lamp through a fiber optic light guide and a diffuser, which provided the intensity and stability required for high speed imaging. A charge coupled device (CCD) camera (Turbo 620G, Stanford Photonics) with custom optics and a resolution of 4.3 µm/pixel at 1000 frames/second, was moved into place over the sample (Fig. 3B). To observe the preparation from the side, we used a 5mm right-angle prism mirror. For all experiments, servo motor position and stimulation timing were preprogramed using custom-made control and data acquisition software written in IGOR (Wavemetrics) running on a desktop PC, and controlled via an A/D board (PCle-6251, National Instruments). Force, motor position, and camera timing pulse were sampled at 20 kHz and saved to a hard disk. Each experimental run was triggered by a signal from the camera corresponding to the first acquired image, which simplified the synchronization of the various signals.

**Figure 3.**
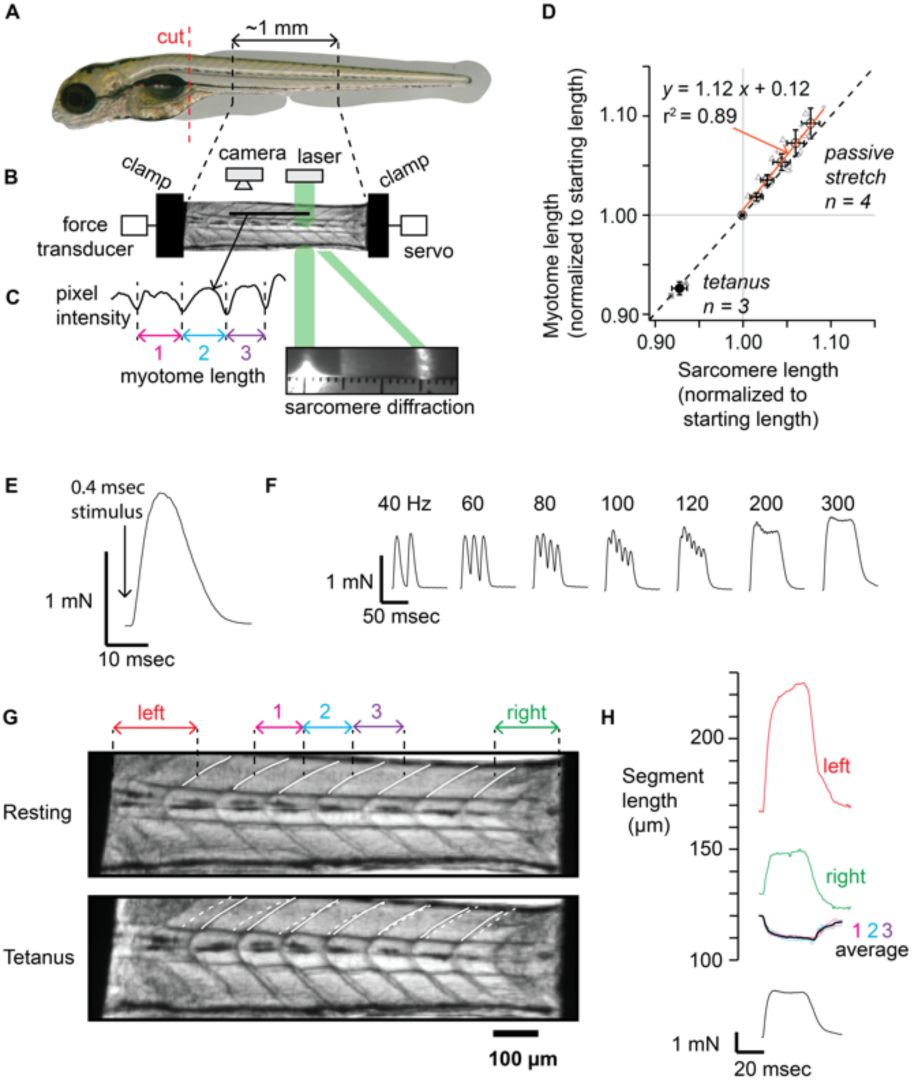
Measurement of force and length in intact zebrafish larval tails. *(A)* A single 5-day old larva as in Fig. 1. Red dashed line represents where larvae are cut prior to mounting for mechanics studies (see Methods). *(B)* An ∼1mm long section of tail in *A* fixed with spring clamps between a force transducer and a servo motor. *(C)* Muscle length in *B* is measured two ways: (1) at the fiber level by automated analysis of 1000 frames/second video images in which a line scan *(black horizontal line in B)* of image pixel intensity detection is analyzed to identify intensity minima, which reflect myotome boundaries (see Methods), or; (2) at the sarcomere level by laser diffraction (see Methods). *(D)* A comparison of proportional changes in myotome vs. sarcomere length in response to passive stretches of preparations to 1.10 x preparation starting length (n=4), and in response to 300 Hz, 150 msec tetanic contractions (n=3). *(E)* The twitch force in response to a single, supramaximal current pulse is complete within 25 msec. *(F)* Increasing stimulus pulse frequencies cause increasing fusion of individual twitch force profiles with a fully fused tetanus at 300 Hz. *(G)* Still frames of a tail preparation from a 1000 frames/second movie (Supplemental Movie 1) prior to *(Resting)*, and during the plateau phase of a 300 Hz, 50 msec tetanic contraction *(Tetanus)*. The position of dorsal myosepta are traced *(solid white lines)* in each image with some used to identify 3 central myotomes and their lengths as in C. To highlight their motion as well as myotomal length changes, myoseptal positions in the Resting image are superimposed on the contracting, Tetanus image (*dashed white lines*). Compliant regions resulting from preparation clamping at both ends are identified as ‘left’ and ‘right.’ *(H)* Longitudinal length change profiles of regions of the tail preparation demarcated by myosepta *(labeled in G)* and clamps throughout the contraction in *G*. The central myotomes *(1, 2, 3, average)* shorten at the expense of compliant regions near the attachment points *(left and right traces).* Tetanic force trace for this contraction at bottom.

After attachment, the length of each preparation was adjusted by moving the position of the force transducer to remove any strain imposed during attachment, and thus returning tail sections to their *in vivo* length. At this point, a single CCD image was used to define the “*starting length”* of the overall preparation and of the central myotomes (described further below). The temperature of the preparation was maintained at 18° C (*length:force, force:velocity*) or 28° C (*isometric force, force:velocity, work loops*) by fresh, oxygenated Ringer’s, pumped at a constant velocity (0.7 ml/min or 6 exchanges/min) through the experimental chamber after being heated or cooled in a custom countercurrent heat exchanger. Tails were then left to rest for 20 minutes before being subjected to one of the experimental protocols detailed below and in the Results.

### Analysis of muscle length during mechanical experiments

Clamping of the tail introduced regions of compliant tissue at each end of the preparation, which allowed the central, functioning myotomes to shorten during contractions by extending the compliant regions. Therefore, we could not use the servo position, i.e. preparation length, as indicative of the myotomal fiber or sarcomere lengths. Diffraction of coherent light by muscle has long been used to determine sarcomere length (25). Therefore, sarcomere length was measured directly using the projected first-order diffraction peak of an ∼200 µm-wide spot of laser light (λ=532 nM) passing through the midpoint of the preparation. A CCD camera recorded the position of the first-order peak, which was projected onto a paper screen (Fig. 3B) so that the distance between the non-diffracted laser beam and the diffracted peak could be converted to sarcomere length using the grating equation. For continuous, high-speed (1000 frames/sec) muscle fiber length measurement, we tracked the three central-most myotomes within the transparent fish (Fig. 3C). Using custom software written in Matlab (Mathworks), we determined the average longitudinal myotome length by first line scanning parallel to the long axis of the tail across 4 myosepta (Fig. 3C). Then the line scan intensity profile was subjected to peak-detection, which identified the high contrast myosepta (i.e. myotome boundaries), with the distance between myosepta equal to myotome and thus fiber length.

### Experimental protocols for tail muscle mechanics

Tail preparations were supramaximally stimulated, using the servo and force transducer clamps as electrodes, with single 0.4 msec current pulses (e. g. *twitches*), or programed trains of 0.4 msec pulses (e. g. *force:frequency, tetanus*) from a biphasic muscle stimulator (MyoPacer, IonOptix).

All experiments involving active muscle contraction were completed within 1 hour of euthanasia. With the exception of the individual twitches evoked during *length:force* experiments, all independently measured contractions were separated by at least 60 seconds, which we found was sufficient to eliminate measurable decline of twitch or tetanus force at a given myotome length within the experimental timeframe. The protocols described below were all accomplished in individual fish.

#### Force:frequency

To establish the stimulation frequency required for a fully fused tetanus at physiologic temperature, preparations maintained at 28°C were stretched to 1.1 x starting length and stimulated for 50 msec with supramaximal 0.4 msec pulses at 40, 60, 80, 100, 120, 200, and 300 Hz. The order of frequencies was alternated between preparations, and a second 300 Hz stimulation was added, either at the beginning or end of the experiment to capture any change in response caused by the intervening contractions.

#### Relationship of myotome length to sarcomere length

To consider myotome length as a proxy for sarcomere length, mounted tails were maintained at 18°C, and either stretched from their starting length to 1.09 x starting length over 2 seconds, or tetanically stimulated (300 Hz, 150 msec.) at their starting length. Since myotome and sarcomere length could not be measured concurrently, each procedure was performed twice. During the first run, preparations were imaged in bright field and myotome length measured as described above. During the second run, sarcomere length was recorded as described above. In both resting and active experiments, the order of myotome and sarcomere length measurements was changed between preparations to account for any systematic differences between the first and second runs.

#### Length:passive force

To assess the contributions of non-muscle structures within the preparation to the passive forces recorded as a function of preparation length, preparations were stretched at varying rates from the starting length to a series of different, as detailed in Results (Fig. 4A, 4B), and the resulting forces recorded. Temperature was held at 18°C throughout the experiment to preserve the integrity of the tail’s attachments to the force transducer and servo. To assess whether these large passive forces compromise our ability to discern the actively generated muscle force, in a control experiment (Fig. 4C, 4D) the preparation was stretched to 1.1 x starting length at 10 lengths/second and held at that length for 6 minutes. The preparation was stimulated with four supramaximal 0.4 msec pulses at 100 millisecond intervals after the stretch, and with the same stimulus pattern five minutes after the stretch. From this experiment, active muscle force for a twitch or tetanus was defined as maximum total force during the contraction minus the force recorded immediately prior to stimulation.

**Figure 4.**
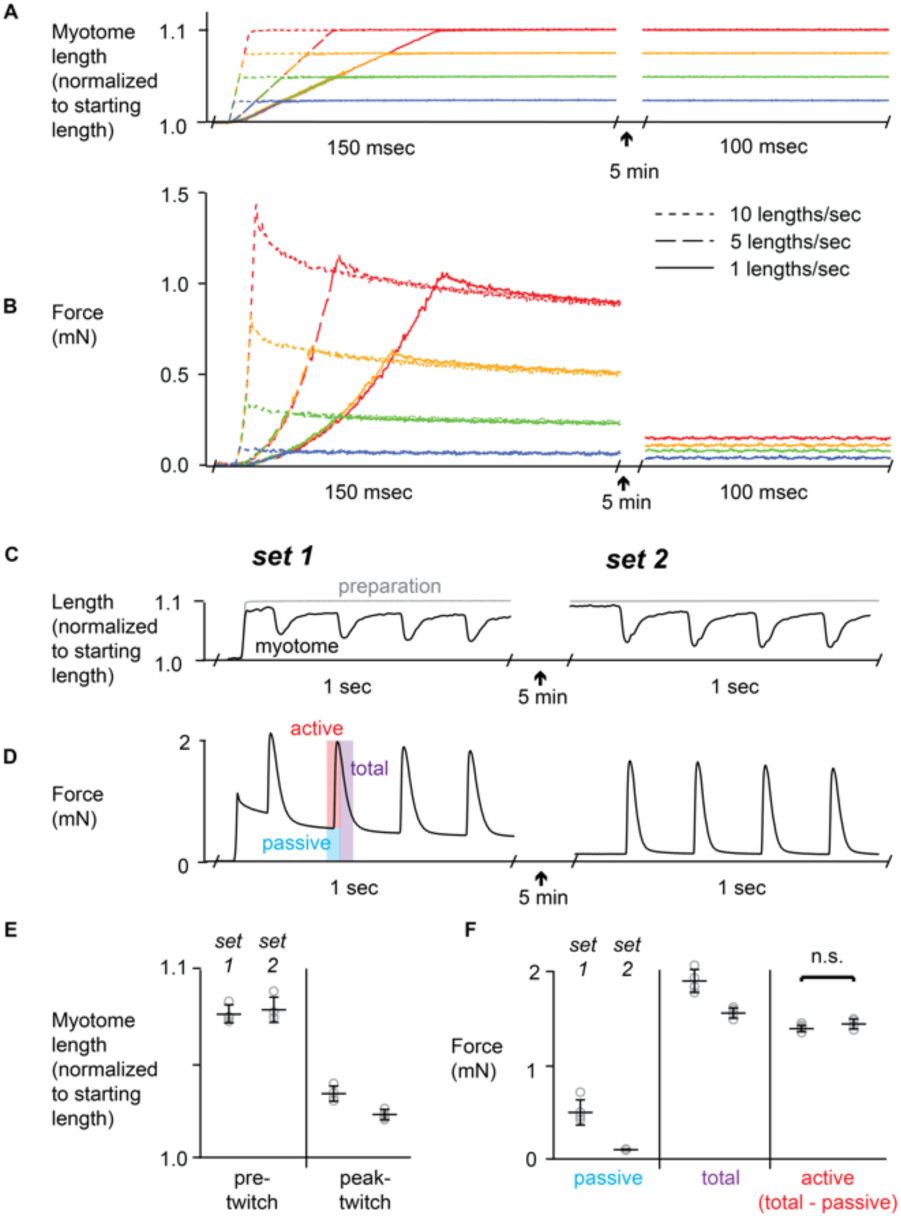
Contribution of passive viscoelastic properties of the tail preparation to measured forces. *(A)* Preparation length traces during servo-imposed stretches to various lengths at various rates. *(B)* Immediate and delayed force responses of a resting tail preparation to stretches in *A*, coordinated by color. The initial, nonlinear response is followed by a decay phase indicative of a slow viscous component not typical of pure muscle tissue. *(C,D)* Preparation length *(gray)* and myotome length *(black)* traces, *C*, and force responses, *D*, of a tail preparation stretched to 1.1 x starting length at 10 lengths/sec (identical to the red, dashed traces in *A, B*), with sets of four twitches immediately (*set 1*), and 5 minutes after (*set 2*), the stretch. *(E)* Myotome length normalized to starting myotome length measured immediately *pre-twitch* and at *peak-twitch* for the two sets of twitches in *F. (F)* Comparisons of passive, active, and total forces as depicted in *D*, between twitch sets. In *E* and *F*, data *(gray open circles)* from individual twitches are shown next to their average ± 1 SD.

#### Length:active force

To relate active force to myotome length, preparations were stretched at a continuous, 0.04 preparation lengths/second for four seconds, with a twitch evoked every 0.5 seconds (Fig. 5A, 5B). Passive force measured just prior to each stimulation was subtracted from the maximum total force of each twitch to calculate active force (Fig. 5B). This approach allowed *length:active force* to be assessed rapidly in a single experimental run, but did not allow preparations to recover after each individual twitch. Therefore, to account for any decline in active force caused by fatigue, the same stimulation pattern was repeated in each preparation without being stretched, as shown in Figure S1. Between the first and last twitch, force declined by an average of 15.5±1.7 %. To correct for this decline in *length:force* measurements, the active force for each twitch evoked during the initial active *length:force* protocol was multiplied by a correction factor calculated from the amount its corresponding twitch declined during the non-stretch control series (Fig. S1).

**Figure 5.**
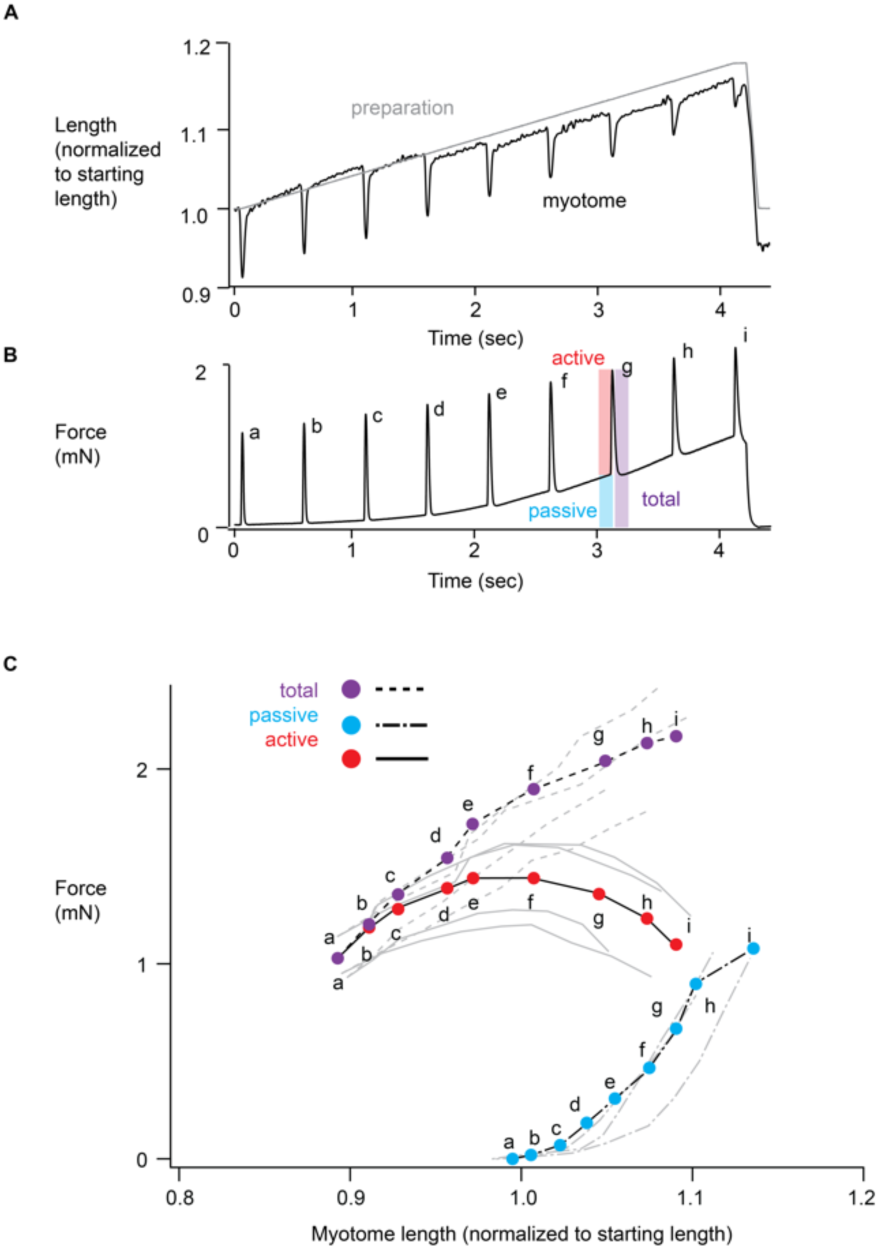
Length:force relationships of larval myotomal muscle. *(A)* Representative preparation length *(gray)* and myotome length *(black)* traces from a tail preparation being stimulated to twitch at 500 msec intervals during a slow stretch to 1.17 x preparation starting length. *(B)* Force trace corresponding to *A*. Passive force is measured immediately prior to the twitch, whereas total force is measured at the peak of each twitch, when myotomes have contracted to their shortest length. Active force is calculated as total minus passive force for each twitch, assuming that passive force remains relatively constant during the twitch (see Fig. 4). *(C)* Forces measured during this experiment for each individual twitch *(denoted a-i)* plotted against normalized myotome length. Data from the tail preparation depicted in *A, B* are highlighted *(colored points)*, while data from 4 additional tail preparations are shown in gray. Total force:length relationships *(dashed lines, purple points)* are shifted to the left relative to passive force:length relationships *(semi-dashed lines, blue points).* The active force:length relationships *(solid lines, red points)* plateaus near the myotome staring length.

#### Isometric twitch and tetanus

Preparations were stretched manually by micrometer to 1.1 x starting length (Fig. 6A). Then a single twitch was followed by 100 msec-long, 300 Hz-tetanus while high-speed movies were recorded and immediately analyzed, as described above, to generate myotome length change profiles (Fig. 6A). In order to maintain myotome length constant (i.e. isometric) during these contractions, we implemented a “feedforward” approach that pre-programed the servo to stretch the preparation during the contraction so as to prevent myotome shortening during the subsequent contractions. Therefore, the myotome length change profiles became the template for programming the servo but in an opposite direction (Fig. 6C). For example, if myotomes shortened by 10% during a contraction, the servo would increase the preparation length 10% over the same time course. Preparations were then shortened by micrometer until myotomes returned to their starting length, and a single twitch and tetanus were evoked while the servo followed the programed stretch profile (Fig. 6C). Finally, the passive force response of the preparation to the stretch profiles were recorded without stimulation (Fig. 6C), and active twitch and tetanus force were calculated by subtracting the passive force traces from the stimulated twitch and tetanus force traces (Fig. 6D). Isometric stress was calculated from force measurements using estimates of muscle cross-section area described below.

**Figure 6.**
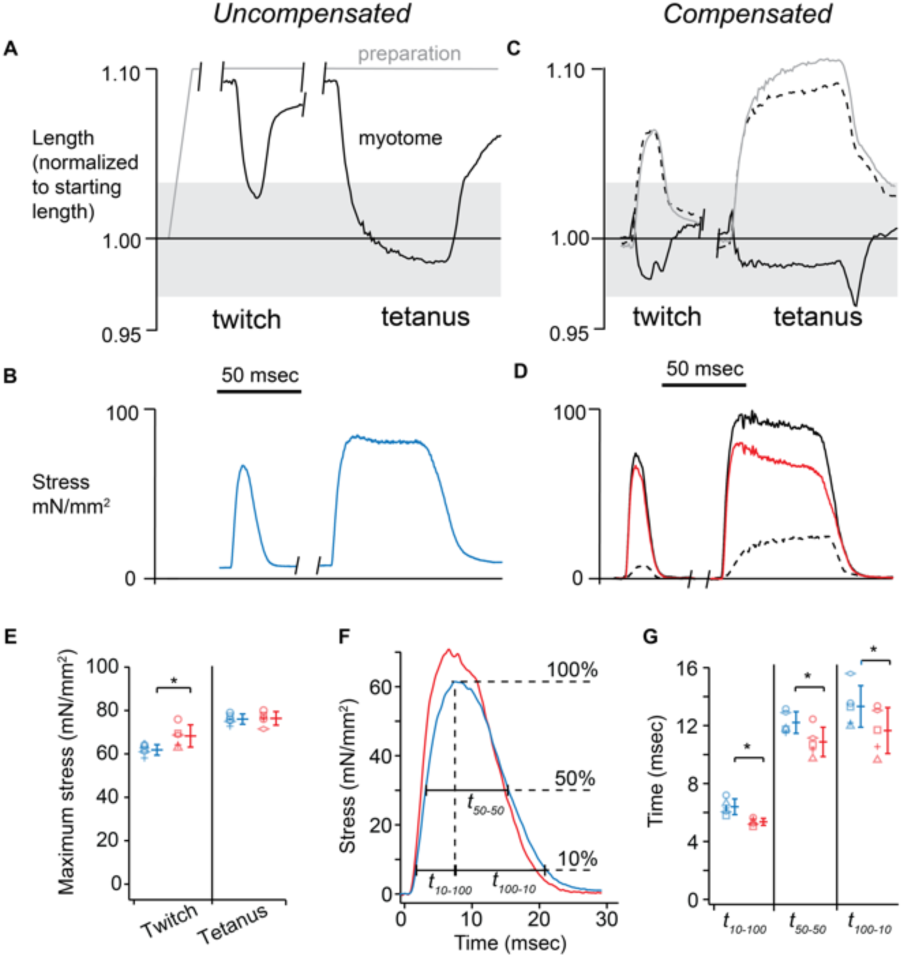
Magnitude and time course of active isometric stress production. *(A)* Representative preparation length *(gray)* and its myotome length *(black)* length traces during a single twitch and tetanus (300 Hz stimulus frequency for 50 msec) several minutes after a stretch to 1.10 x starting myotome length. *(B)* Stress traces (force/muscle CSA – see Methods) corresponding to *A. (C)* Myotome length profile in *A* is used to program a compensatory “feedforward” stretch (*gray trace*) in subsequent contractions of the same preparation during a twitch and tetanus. By applying these corrective stretches during the time course of the contractions, the myotome lengths remained near-isometric *(solid black trace)*. The dashed black trace shows myotome length in response to the compensatory stretch without the preparation being stimulated. Shaded regions in *A* and *C* represent starting myotome length± 3% - the range over which maximum active force is developed. *(D)* Stress traces corresponding to *C*: stimulated (solid black trace) and unstimulated (dashed black trace). Active stress *(red trace)* is calculated as the difference between stimulated and unstimulated stress traces. *(E)* Maximum stress during uncompensated (*blue*) and compensated (*red*) twitches and tetani at 28°C (n=5 tails). *(F)* Representative uncompensated *(blue trace)* and compensated *(red trace)* twitch stress traces. *(G)* Time courses - as depicted in *F* - for stress development *(t*_*10- 100*_), full width at half-maximum stress *(t*_*50-50*_), and relaxation *(t*_*100-10*_) in uncompensated *(blue)* and compensated (*red*) twitches at 28°C (n=5 tails). Data from individual tails *(open symbols)* are shown next to their average ±1 SD (*P < 0.05).

#### Force:velocity

To measure the effect of shortening velocity on active force during steady state activation, preparations were initially stretched to 1.08 x starting length by micrometer, followed by a servo-based stretch at 0.5 preparation lengths/second to 1.12 x starting length (Fig. 7A), before being tetanized (100 msec, 300 Hz) (Fig. 7C). At 40 msec after the initiation of the tetanus, preparations were shortened by 4% (i.e. 1.12 to 1.08 x starting length) at a constant velocity (Fig. 7A). Ramps were limited to 4% of preparation length to preserve preparation integrity. Under these conditions, we were limited to preparation shortening velocities less than 7 lengths/sec by the resolution of the equipment. The same protocol was then repeated without stimulation in order to measure the tail’s passive force response to the length changes. The passive force trace was subtracted from the stimulated force trace to determine active force. The proportion of maximal active isometric force developed while shortening at a particular velocity was defined as the minimum force achieved prior to the end of the shortening ramp divided by the active force measured 20 msec after the end of the ramp, at which point the recovery of isometric force had reached a plateau. Myotome length was measured throughout the experiment as described above, and myotome shortening velocity was calculated as the proportional change in myotome length over the duration of the ramp. Stimulated and passive shortening runs were repeated in the same preparation at multiple preparation shortening velocities: 1, 1.5, 2, 3, 4, 6 preparation lengths/second at 18°C, and; 1.5, 3, 6 preparation lengths/second at 28°C.

**Figure 7.**
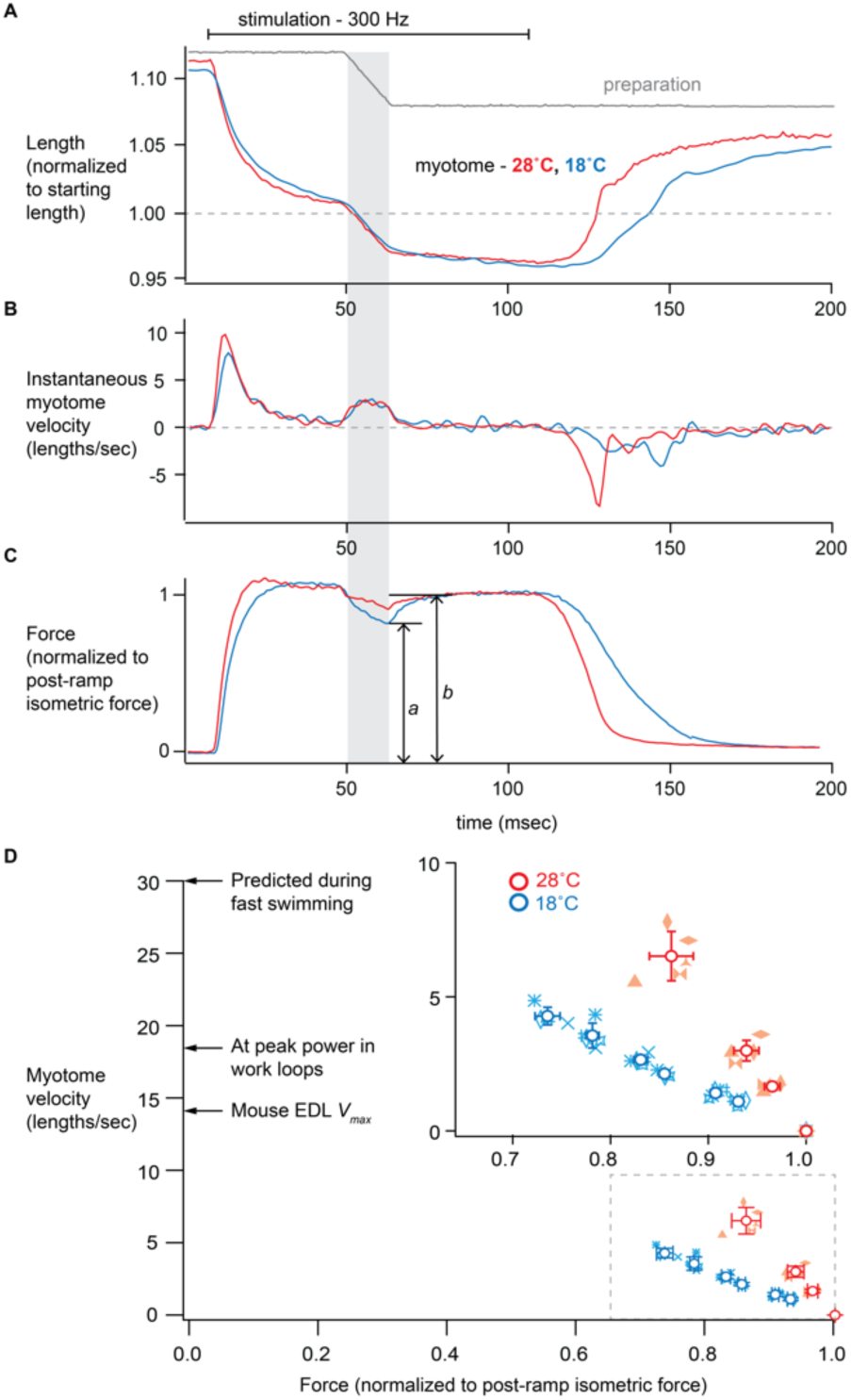
The effect of shortening velocity on steady-state active force. *(A)* Representative preparation *(gray)* and its myotome length *(blue – 18°C, red – 28°C)* traces during isovelocity shortening ramps imposed mid-tetanus (300 Hz, 100 msec) in tails pre-stretched to 1.12 x preparation starting length. *(B)* Instantaneous velocity and *(C)* active force traces normalized to forces to which the preparations recover (*b*), post-shortening ramp, as imposed in *A*. The shaded vertical bar represents the period of the isovelocity ramp. Active force associated with a particular myotome shortening velocity was that at the end of the isovelocity ramp to the post-ramp recovered isometric force, *a* (see Methods). *(D and inset)* Relationship of active force to measured myotome shortening velocity from the experiment depicted in *A, B, C* at multiple velocities (n= 5 tails at each temperature). Individual data points from each tail are shown *(symbols)* along with average ± 1SD data for each imposed velocity. The large graph shows data from inset (within gray dashed box) in the context of inferred *in vivo* larval muscle shortening velocities during fast swimming (30 lengths/second, 28°C) (26), V_max_ of mouse EDL (∼14 lengths/second, 37°C) (45), and myotome shortening velocity observed at peak instantaneous power in 100 Hz work loops (18.1±2.1 lengths/second, 28°C) as described below and in Figure 8.

#### Cyclical work and power

For work loop protocols, the preparation was initially stretched by micrometer to 1.05 x starting length. This value was chosen because it resulted in myotome lengths being within ± 5% of starting length during the work loops. A work loop was achieved by sinusoidally varying the myotome length between 1.01 and 1.09 x starting length for ten cycles at a fixed frequency. We chose this 8% maximum strain since it did not lead to rapid deterioration of preparation at its attachment points and was near to the predicted 10% *in vivo* strain for fast fibers (26). Prior to the shortening phase of each of the first five cycles, preparations were stimulated with a single supramaximal 0.4 msec twitch pulse. The phasing of this pulse was set so as to maximize net positive work for each cycling frequency (27). The protocol was repeated in each preparation at 40, 60, 80, 100, and 120 Hz cycling frequencies. Work and power measurements were made using averaged data from the second, third, and fourth stimulated loops at each frequency. Particular care was taken to synchronize myotome length and force signals using the camera timing pulse, since even a small offset at the higher frequencies would result in large changes in estimated work and power. As in other protocols, the order of frequencies was reversed in half of the preparations studied.

### Data analysis

To calculate active muscle stress (mN/mm^2^) we divided active force (mN) by 2.18×10^−2^ mm^2^, which was the cross-sectional area (CSA) of muscle measured at the anal fin in histological sections (described above). This was possible due to the consistency of size among larvae studied. For example, the dorsal-ventral height measured at the anal fin in video images varied between individual preparations by less than 5% (238±7 µm, n=13), which was not significantly different than the equivalent measurement made in histological sections (236±5 µm, n=3). For muscle-weight-specific work and power, we estimated the muscle weight by assuming the volume of muscle within a central myotome was equal to the muscle CSA times the starting myotome length, as measured above, times a density of 1.06 g/cm^3^ (28). Work per cycle, peak power and net power produced by this volume were calculated as in (27). Myotome length and myotome velocity for these calculations were taken directly from myotome length analysis described above.

### Statistical analysis

Data are reported as mean ± 1 standard deviation. Dependent variables were compared using two-tailed, paired t-tests when treatments occurred within individual preparations, or unpaired t-tests where not. Linear regression was used to describe the relationship between sarcomere and myotome length in passively stretched tails. Statistical calculations were performed using IGOR (WaveMetrics) software.

## Results

### Anatomical features of the 5-day old zebrafish tail

The usefulness of the larval zebrafish tail as a model system to probe muscle physiology and mechanics depends on the ability to translate length and force measurements from the intact larvae to their origins in the locomotory muscle fibers and their sarcomeres. Therefore, to define the gross morphology of the tail as it relates to the arrangement of muscle fibers and other non-muscle structures, we used live imaging, together with light and transmission electron microscopy (TEM) of fixed sectioned tissue.

Figure 1 shows major anatomical features of the tail from a 5-day post-fertilization larva in longitudinal (Fig. 1A, 1B) and transverse (Fig. 1D, 1E) views. The locomotory musculature runs nearly the full length of the tail and consists of a series of ∼25 myotomes in a chevron pattern that are separated by connective tissue boundaries, i.e. myosepta (Fig. 1A-1C). Individual myotomes consist of bundles of striated muscle fibers (100 µm in length) that span the myotome and insert into myosepta at both ends (Fig. 1B), with fibers oriented nearly parallel to the long axis of the tail (16). In the tail’s central region, the locomotory muscles occupy 67.3±0.4% of the larva’s cross sectional area (CSA), and surround both the spinal cord and notochord, which is a semi-rigid connective-tissue sheath that surrounds cells with large pressurized vacuoles (29, 30) (Fig. 1D, 1E). Both the spinal cord and notochord run the full length of the tail. Both fast- and slow-twitch muscle fibers are expressed at this early stage of development and are anatomically segregated (23). Specifically, the slow-twitch fibers occupy a thin, single-cell layer near the tail’s lateral surface, making up 5.4±0.2% of the tail CSA (Fig. 1E). Whereas, fast-twitch fibers are dominant, constituting 61.2±0.5% of the tail CSA (Fig. 1E).

The ultrastructural appearance of fast-twitch muscle fibers is typical of vertebrate striated muscle as seen in longitudinal (Fig. 2A, 2C), and transverse (Fig. 2B, 2D) TEM images. Specifically, individual fibers contain myofibrils, having clearly defined sarcomeres (1.84±0.03 µm long) with identifiable Z- and M-lines, as well as A-bands representing regions of overlap of the thin (actin-containing) and thick (myosin-containing) filaments (Fig. 2C). Myofibrils are surrounded by a well-developed sarcoplasmic reticular membrane network with enlargements of this network (i.e., lateral sacs) near the Z-lines that are juxtaposed to the transverse tubules (Fig. 2A-D). In cross section, the highly ordered hexagonal lattice of thin filaments surrounding each thick filament is apparent (Fig. 2D, inset).

### Measuring force and length in intact, mounted tail sections

To measure the force and shortening generated by the longitudinal, fast-twitch muscles within the tail, we clamped ∼1 mm sections of 5-day old larval tails at their *in vivo* (starting) length between a force transducer and high-speed servo, length controller (Fig. 3A, 3B). Because tails are transparent, we were able to measure sarcomere length directly in mounted tails by laser diffraction, which at their starting length (1.87 ± 0.02 µm) were similar to histological measurements as reported above. Since the quality of the laser diffraction signal was highly sensitive to position and motion, sarcomere length could not be monitored reliably during the time course of active muscle force development. Therefore, we investigated whether myotome length, as measured along the long axis of the tail, could be used as a proxy for muscle fiber and sarcomere length during mechanical experiments. For this we developed custom Matlab software to identify and track myosepta, which appear in brightfield video images as clearly defined dark chevron lines (Fig. 1A). Specifically, the software uses peak-detection to measure the distances between adjacent myosepta (i.e. myotome length) along a scan line drawn parallel to the long axis of the tail (Fig. 3B, 3C). When tails were stretched while relaxed between 1.00 and 1.09 x starting length, myotomes and sarcomeres measured near the center of the preparation, increased in length by a similar amount (1.09 ± 0.02 and 1.08 ± 0.01 x starting length, respectively, n= 4) (Fig. 3D).

When held at a fixed length and stimulated with a sub-millisecond current pulse directly through the clamps at 28°C (see Methods), tails responded with a characteristic ‘twitch’, where force rose rapidly and returned to baseline within 25 msec after each stimulation (Fig. 3E). Similar twitch responses were observed at stimulation frequencies up to 40 Hz (Fig. 3F). At higher frequencies, individual twitches merged, forming fully fused tetani at 300 Hz (Fig. 3F). The short twitch duration and high tetanus stimulation frequency were consistent with the very high tail beat frequencies observed *in vivo* (26, 31). However, even though the tail length was held constant (i.e., isometric) during this stimulus protocol, myotomes consistently changed length during contractions, as measured in movies taken at 1000 frames/second (Supplemental Movie 1). Specifically, for either a twitch or tetanus contraction, interior myotomes, i.e. those not in immediate proximity to the clamps, shortened along the long axis of the tail by as much as 15% in phase with force generation, while myotomes near the ends of the preparation lengthened (Fig. 3G, 3H). We interpreted this strain pattern to be the result of local tissue damage caused by the crushing action of the clamps themselves. Specifically, during contraction, the damaged and thus compliant regions at the ends of the preparation are distended by the forces developed by the central healthy myotomes. This conclusion was supported by the visual appearance of the tail section, since myotomes that shortened during contractions also had clearly defined myosepta, whereas those near the clamps did not. To standardize our approach, and to avoid the boundaries between healthy and damaged regions of the preparation, we defined myotome length as the average length of the three central-most adjacent myotomes (Fig. 3B, 3C, 3G, 3H). Steady state sarcomere length measurements, from the central part of the preparation, prior to stimulation and during the tetanus plateau also showed that sarcomeres shortened by the same proportion as did myotome length measured in this manner (Fig. 3D, n=3). This observation validated that myotome length was a good correlate for muscle fiber and sarcomere length during the various experimental protocols to be described and was used throughout this study both under relaxed and activated tail conditions.

### Viscoelastic properties of the resting tail preparation

The relationship between muscle length and force under both relaxed (i.e. passive) and active conditions are muscle’s most basic physiological characteristics (32). Since fast muscle fibers run nearly parallel to the long axis of the tail (16) (Fig. 1), we can probe these relationships by changing the preparation length. However, the structural complexity of the tail, with its sizeable notochord and spinal cord, may contribute to the passive viscoelastic properties of the tail preparation as these structures are stretched. To characterize any such viscoelastic contributions, we stretched a non-stimulated preparation at constant velocities (1-10 lengths/sec) to a series of lengths longer (2.5-10%) than the starting length and then measured the resulting force response over time (Fig. 4A, 4B). Increasing preparation length resulted in an initial, nonlinear rise in force coincident with the length change, followed by a relaxation in force over several minutes to a much lower steady-state level at the new, longer length (Fig. 4B). The initial nonlinear force:length relationship is typical of most biological tissue (e.g. tendon (33), lung (34)) including muscle (35). Unlike muscle, however, the majority of the tail’s force-response to stretch decays with time, reflecting the viscoelastic nature of structures within the tail (e.g. notochord). Therefore, under activating conditions, the contributions of these viscoelastic elements to the total force generated by the tail must be accounted for in order to characterize the underlying muscle fibers’ active force generation. However, upon stimulation, myotomes shorten (Fig. 3G, 3H). When doing so, do the myotome length changes impact the length of the structures responsible for the passive viscoelastic forces described above, or do these structures maintain their length and with it their contribution to the total force?

To distinguish between the two possible scenarios described above, we evoked a series of twitches immediately after a 10 length/sec stretch to 1.1 x starting length, and 5 minutes later when the passive force at the longer length had decayed to its low steady-state level (Fig. 4C, 4D). For each twitch, the passive force prior to the twitch and the total force at the peak of each twitch were recorded, which allowed active force to be defined as total minus passive force (Fig. 4D, 4F). In addition, the maximum myotome shortening at the peak of the twitch was determined (Fig. 4C, 4E). Interestingly, the active twitch force was constant, and myotome shortening similar, regardless of the time following the stretch (Fig. 4E, 4F). These results suggest that differences in total twitch force between the two time points reflects the degree to which passive force decays, with the extent of this force decay similar for protocols with and without superimposed twitches (Fig. 4B, 4D). More importantly, the time-dependent component of the tail’s viscoelasticity is unaffected by the change in myotome length that accompanies contraction and thus, active force is a true measure of the underlying tail muscle.

### Length dependence of active force

Having established the passive viscoelastic force:length properties of the relaxed tail, we determined the dependence of active force on myotome length. Therefore, tail preparations (n=5) were stretched from 1.00 to 1.17 x starting length over 5 seconds, while evoking twitches every 0.5 seconds (Fig. 5A, 5B). Myotomes shortened during each twitch with the extent of shortening diminishing at the longer preparation lengths (Fig. 5A). To relate the active force generation to the myotome length at the peak of each twitch, we defined active force, as above, as total force minus passive force measured at the same preparation length. However, we plotted the total and active force versus the myotome length achieved at the peak of the twitch (Fig. 5C). By this protocol, the range of myotome lengths during active contractions varied between 0.90 and 1.10 x starting myotome length. Both passive and total twitch force increased with increasing preparation and myotome length, whereas active twitch force reached a peak near 1.00 x starting myotome length and decreased at myotome lengths longer than their starting length (Fig. 5C). Interestingly, over the range of myotome lengths between 0.95 and 1.05 x starting myotome length, more than 90% of the peak active twitch force could be generated; equivalent to the range of lengths predicted to be utilized *in vivo* (26).

### Maximum twitch and tetanic stress under near-isometric conditions

The time course and magnitude of active myotome stress (active force/muscle CSA) generation are dependent on both stimulus frequency (Fig. 3F) and myotome length (Fig. 5C). Thus, with myotomes shortening by between 5 and 15 % during a twitch and tetanus (Fig. 3G, 3H, 4E, 5A), the maximum active, isometric force generating capacity at the myotome’s optimal length could not be determined. Given that myotome shortening occurred within the first 10-15 milliseconds after stimulation, real-time length feedback based on video image analysis was not possible due to processing time. Therefore, we used an iterative feedforward approach to compensate for myotomal motion during twitch and tetanus contractions (Fig. 6).

To maintain the preparation isometric at the myotome level during the course of a contraction, we recorded the time course of myotome shortening following a stimulus (Fig. 6A, 6B) and then, in a subsequent contraction, applied a signal to the servo length motor that would stretch the preparation in an equal and opposite manner. In order to compare both twitch and tetanus stress before and after myotome length compensation at the myotome’s optimal length for stress production (Fig. 5C), it was necessary to first pre-stretch tails (n=5) to 1.10 x starting length (Fig. 6A) so that when myotomes shorten during a twitch and tetanus, their final shortened lengths were within ±3% of the myotome’s starting length (i.e. the optimal length for active stress production, Fig. 5C). The myotome shortening response was recorded, as described above, and used as the effective feedforward signal to the length servo. Next, the preparation length was returned to its starting length and a twitch and tetanus evoked but with the compensatory length control applied (Fig. 6C, solid gray traces, Supplemental Movie 2). Using this protocol, myotome length was maintained near-isometric with at most a 3% deviation from the starting myotome length prior to the stimuli (Fig. 6C, solid black traces). Finally, the correction-stretch protocol was repeated without stimulation (Fig. 6C, 6D, dashed traces), and the passive stress responses subtracted from the length controlled, total twitch and tetanus stress responses in order to determine corrected active stresses (Fig. 6D, red trace). Reducing myotome shortening by this method had its greatest effects on both the magnitude (Fig. 6E) and the time course of active twitch stress (Fig. 6F, 6G) with no effect on active tetanic stress (Fig. 6E). Specifically, active twitch stress increased significantly by 10% when maintained near-isometric (Fig. 6E) with the activation time *(t*_*10-100*_), full width at half-maximum stress *(t*_*50-50*_), and relaxation time *(t*_*100-10*_), significantly shorter (Fig. 6G). Using compensated values, the twitch to tetanus ratio of maximum stress was 0.91±0.05 (n=5).

### The effect of shortening velocity on active force

The force versus velocity relation is another of muscle’s most basic physiological characteristics, where active force decreases with shortening velocity (36). Therefore, to define this relationship in the larval tail at 28°C, we applied 4% shortening ramps at the plateau of tetanic contractions at velocities between 1-7 myotome lengths/sec (Fig. 7A). Prior to contractions, tails were stretched (see Methods) so that during force generation myotomes would shorten to their near optimal length for peak active force before the shortening ramps were applied (Fig. 7A). In response to these ramps, force declined and the value at the end of the ramp (*‘a’* in Fig. 7C) was normalized to the force that the tail recovers to isometrically at the new shorter length (*‘b’* in Fig. 7C). In response to the shortening ramps, normalized force decreased with increasing ramp velocity (Fig. 7D). Interestingly, myotomes could maintain 0.86±0.02 of their isometric force levels while shortening at 6.5±0.9 myotome lengths/sec. To potentially extend the range of the force versus velocity relation, we reduced the experimental chamber temperature to 18°C. Even at this temperature, myotomes maintained 0.74±0.1 of their isometric force when shortening at 4.3±0.3 myotome lengths/sec (Fig. 7D). Lowering temperature by 10°C also reduced the peak instantaneous myotome shortening velocity achieved during force development following the stimulus from 13.2±2.1 lengths/sec. to 9.7±1.1 lengths/sec. (Fig. 7B, 7D).

### Cyclical power production at approximate in vivo strains and frequencies

To power larval swimming, the tail musculature must undergo repetitive cycles of sequential activation, force production while shortening, and lengthening by opposing muscles during relaxation. Power during a complete locomotory cycle is defined as the net positive work per cycle time, with net work determined as the work produced during shortening minus the work absorbed during lengthening. At 5 days, zebrafish larvae swim with tail beat frequencies up to 100 Hz (26, 31) and undergo estimated muscle strains of 10% (±5% of resting tail length) (26). To measure the capacity of myotomal muscle to produce cyclic power under approximate *in vivo* strains and cycle frequencies, we used a modified work loop approach (37). Specifically, preparation length was varied by 8% sinusoidally at 40, 60, 80, 100, and 120 Hz at 28°C (Fig. 8A). The tail preparations (n=5) were stimulated with a single pulse at a time in the sinusoidal length-cycle that maximized net work in the active work loop (Fig. 8A, 8B). Passive work loops were also obtained in relaxed preparations (Fig. 8A, 8B). Myotome lengths were measured at every point in the cycle so that work loops could be generated by plotting force versus myotome length at every time point (Fig. 8B).

**Figure 8.**
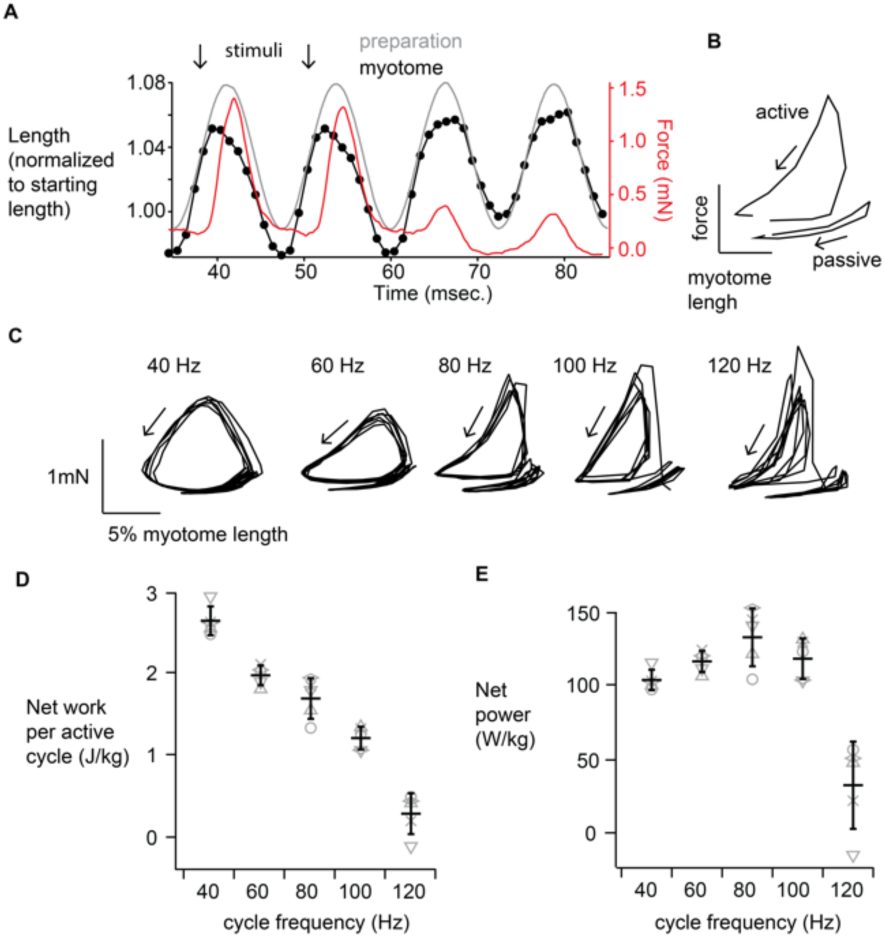
Work and power production by tails under simulated *in vivo* contraction regimes. *(A)* Representative work loop length and force traces in the time domain. An imposed, 80 Hz sinusoidal preparation length change *(gray trace)*, results in a similar change in myotome length *(black data points)*. During each of the first two cycles, the tail is stimulated (black arrows) to twitch such that force *(red trace)* is highest during the shortening phase. The second two cycles are unstimulated. *(B)* A representative work loop cycle with force as a function of myotome length. Net work done during the active work loop cycle is proportional to the area within the counterclockwise loop. During the passive work loop cycle, a small amount of work is absorbed and is proportional to the area within the clockwise loop. *(C)* Representative work loops (5 active above, followed by 5 passive below, as in *B*) at each of the cycling frequencies studied. *(D)* Net work per active work loop cycle as a function of cycle frequency (n=5 tails). *(E)* Net power per active work loop cycle as a function of cycle frequency (n=5 tails). Data in *D* and *E* from individual tails (*open symbols*) are shown with their average ± 1 SD.

Figure 8B shows a representative 80 Hz active work loop followed by passive work loop. For the active work loop, the loop trace goes counterclockwise, which represents positive net work production estimated by the area within the loop. Whereas, in passive work loops, the loop traces are collapsed and clockwise, representing very low net negative work (i.e. the preparation absorbs work from the servo). Net work per active work loop decreased with increasing cycle frequency from 2.63±0.18 J/kg of muscle at 40 Hz to 0.43±0.27 J/kg at 120Hz, (Fig. 8C, 8D). However, maximum net power (Fig. 8D) was produced at 80 Hz (134.4±19.9 W/kg) and remained high (122.9±11.1 W/kg) at 100 Hz, which is the maximum tail-beat frequency observed during fast larval swimming (21), before falling off steeply at 120 Hz. Peak power during the shortening phase was 498.9 ± 127.0 W/kg (80 Hz) and 640.2 ± 151.5 W/kg (100 Hz). These numbers exceed by approximately 50% the estimates of muscle power density required for this mode of swimming based on a kinematic analysis of tail motion (20). Interestingly, peak power was generated by the active preparations while shortening at 18.1 ± 2.1 myotome lengths/sec during 100 Hz work loops. (Fig. 8A).

## Discussion

The advantage of using larval zebrafish for studying muscle biology is based on the relative ease with which this model system allows genetic and other manipulations to be introduced in a fully functioning muscular system, coupled with the ease of measuring the effect of such experimental interventions on muscle mechanics. Remarkably, as demonstrated here and elsewhere (4, 8, 17, 18), larval tails can be mounted without further dissection in a muscle testing apparatus so that forces developed during contractions are measured along the long axis of the tail. We describe two significant improvements to this general approach by correcting for: 1) the significant mechanical contribution of non-muscle structures (i.e. spinal cord, notochord), and; 2) the internal myotomal shortening that occurs during isometric tail contractions. These corrections allowed us to draw inferences from tail mechanics about force and motion generation at the level of the sarcomere. Our results suggest that myotomal muscles are designed to operate at extreme speeds and power necessary to support bursts of larval swimming *in vivo.*

### Isometric twitch and tetanus

Myotomal muscle’s rapid twitch and the high stimulation frequency (300 Hz) to achieve a fully fused tetanus are both consistent with larvae’s ability to swim with tail-beat frequencies approaching 100 Hz (26). In vertebrate muscle, the time courses of force development and relaxation depend in part on the rates of Ca^2+^ delivery to and removal from the cytosol in response to an excitatory stimulus. The ultrastructural anatomy of myotomal muscle (Fig. 2) reveals adaptations to the sarcoplasmic reticulum that would speed the rates of these calcium handling processes: a) highly enlarged lateral sacs, the site of myoplasmic Ca^2+^ release; b) an extensive and well distributed sarcoplasmic reticular network surrounding each myofibril that contains SERCA pumps to sequester Ca^2+^ back into the sarcoplasmic reticulum during relaxation (38). The fact that twitch-forces nearly equal that during tetani suggests that Ca^2+^ released upon activation must reach near-saturating levels within the time course of a twitch.

### Length versus force relationship

The dependence of active muscle force generation on muscle length is one of muscle’s most basic physiological properties and is set by the overlap of the interdigitating thick, myosin and thin, actin filaments within the sarcomere (Fig. 2). Classically, the optimal sarcomere length for active force generation occurs when overlap of the thick and thin filaments allows for the maximum number of attached force-generating myosin crossbridges to the thin filament (32). At sarcomere lengths both shorter and longer than this optimum, active force generation is reduced. Interestingly, most skeletal muscles *in vivo* take advantage of the length:force relationship by operating near or at their optimal sarcomere length (39, 40). In previous larval tail mechanical studies (8, 17, 19), in which sarcomere length was measured prior to each contraction, maximal stimulated forces were recorded at lengths 8-12% longer than the *in vivo* resting length. This finding suggested that larval swimming would be powered by muscles operating at sarcomere lengths well below their optimal range for force development (26). More likely is that significant internal fiber shortening may have occurred in these earlier studies due to preparation compliance as reported here. Once corrected for this internal shortening (Figs. 3, 6), the optimal length for maximal myotomal force production (Fig. 5) does coincide with the larva’s *in vivo* resting tail length, where it most likely operates during swimming.

### Shortening velocity and power generation

In 1938, A.V. Hill described the now classic hyperbolic relationship between a muscle’s shortening velocity and the load against which it is shortening (36). At the extremes, a muscle shortens at its maximum velocity (*V*_*max*_) under unloaded conditions and does not shorten under isometric conditions, where the muscle generates its maximum force. Although muscles do not generate power (i.e. force x velocity) at these two extremes, muscles do produce their maximum mechanical power when shortening in the range between 0.15 to 0.40 *V*_*max*_, and in general locomotory muscles are biomechanically designed to operate within this range *in vivo* (41-43). Given that the tail muscles power the larva’s fast swimming movements, we sought both to characterize their force versus velocity relationship and to estimate power production through a work loop protocol, which would allow us to compare these physiological parameters to estimates derived from larval body kinematics during swimming (26).

To describe the force:velocity relationship, we imposed length changes to the tail at known velocities during the force plateau of a tetanus and then measured the resultant reduction in force (Fig. 7). However, the temporal resolution of our instrumentation limited the speed of the length changes to a maximum of 6.5 myotome lengths/sec. Interestingly, this speed only reduced the active tetanic force by 14% at 28°C. Thus, we were unable to fully characterize the shape of the force:velocity relationship over its entire range of forces and thus, would require instrumentation that could impose and record the presumably high speeds of shortening that these larval tails are capable of (Fig. 7). In fact, body kinematics during swimming predicted that fast myotomal muscle fibers in the tails of 5-day old larvae shorten at up to 30 lengths/sec (Muller, 2004), which is not even *V*_*max*_ since swimming is a loaded movement. Based on our high-speed myotome imaging during periods of force development following a stimulus, we observed internal myotomal shortening velocities of greater than 13 lengths/sec at 28°C (Fig. 3H, 7B, 7D), which once again is not *V*_*max*_. Presumably, these high shortening velocities, regardless of their exact absolute values, reflect the inherent actomyosin ATPase activity of the fast-skeletal myosin heavy chains that are expressed in the developing larvae (44).

Without a complete force:velocity relationship to provide an estimate of peak power production and the velocity at which this occurs, we used a work loop protocol to estimate work and power directly under approximate *in vivo* conditions. Interestingly, the larval tails produced high net power over the entire range of their *in vivo* work loop frequencies (Fig. 8) (26). More amazing is that during this protocol, peak power was highest at a work loop frequency of 100 Hz, when myotomes were shortening at more than 18 lengths/sec (Fig. 8A). Therefore, if we assume that this velocity represents 0.4 *V*_*max*_ (see above), we can make a rough, but conservative estimate of myotomal *V*_*max*_ to be 45 lengths/sec at 28°C; which would make it the fastest shortening vertebrate locomotory muscle by a factor of ∼2 (Medler, 2002). As a benchmark, the well-studied, predominantly fast-twitch mouse extensor digitorum longus (EDL) at 37°C has a reported *V*_*max*_ of ∼14 muscle lengths/sec (45).

These larval myotomal muscles in certain ways resemble the ‘superfast’ vertebrate muscles, found exclusively in sound-producing organ systems, which work at cycling frequencies between 100 and 200 Hz (27, 46-48). The physiologic adaptations that enable superfast muscle to cycle at such high frequencies come at the expense of force production. Specifically, a large proportion of the muscle cell volume and energy usage are dedicated to calcium handling (Fig. 2), which reduces available space for sarcomeres. Also, to speed relaxation, superfast muscles express myosins with high rates of detachment from actin, resulting in a low duty-cycle molecular motor with low force generation (48, 49). Myotomal muscle may make similar trade-offs: at 76 mN/mm^2^ (Fig. 6), myotomal muscles generate maximum isometric stress intermediate between mouse EDL (243 mN/mm^2^) (45) and the ‘superfast’ swimbladder muscle of the oyster toadfish (24 mN/mm^2^) (49). Where myotomal muscle is unique is in its power generating capacity. Despite the zebrafish larvae developing moderately low active isometric stresses, their unique force:velocity characteristics mean they produce muscle-weight specific power in sinusoidal work loops similar to mouse EDL, but at ten-fold higher cycling frequencies (122 W/kg @ 100 Hz vs. 132 W/kg @ 9 Hz, respectively) (45).

How does the extreme mechanical performance of larval zebrafish muscle impact its value as a model system? While these muscles are not as close mechanically or genetically to human muscle as those of the mouse, they present a unique opportunity to address fundamental questions. For example, how are vertebrate muscle-mechanical properties like *V*_*max*_ tuned to match the constraints imposed by specific muscle-powered behaviors? While zebrafish require ultrafast muscle performance in order to swim as larvae, the biomechanical and environmental demands on their locomotory apparatus change continually as their mass increases by 1000-fold during growth to adulthood (21). How this is accomplished at the sarcomere level is likely relevant to human physiology, since those genes (e.g. myosin binding protein C) known to regulate the muscle mechanical relationships considered in this study have homologs in zebrafish (4).

## Conclusion

Larval zebrafish have potential as an experimentally flexible model system to study muscle function in health and disease due to their rapid development and genetic tractability. The architecture of the larval locomotory musculature allows intact tails to be studied using the equipment and approaches developed to study *ex vivo* muscle mechanics. Refinements to this approach described in this study, which incorporate accurate measurements of muscle length, and active force, enable the application of a suite of classical muscle-mechanics experiments to define fundamental contractile parameters of the muscle. With this approach in hand, the larval tail becomes an intact muscle mechanics preparation that is amenable to high-throughput drug screening or highly iterative transgenics that would be impossible or cost prohibitive in the mouse. In addition to being useful as a model system, zebrafish larval muscle is shown to function at shortening velocities exceeding any known vertebrate skeletal muscle. This observation alone warrants further study to determine the molecular adaptations within the sarcomere that enable such extreme mechanical performance.

## Author Contributions

A.F.M. designed research, performed research, contributed analytic tools, analyzed data, wrote the manuscript. G.G.K. designed research, contributed analytic tools. B.M.P. designed research, contributed analytic tools. A.M.E. designed research. D.M.W designed research, contributed analytic tools, wrote the manuscript.

## Acknowledgements

The authors thank Marion Siegman for donating the muscle mechanics apparatus without which none of the experiments would have been possible, Ashley Waldron for expertise with the zebrafish model, Samantha Previs for technical support, and Shane Nelson for analytical advice, as well as Michele von Turkovich and Douglas Taatjes from the Microscopy Imaging Center for morphological expertise.

This work was supported by funds to D.M.W. from the National Institutes of Health (AR067279) and a generous gift from Arnold and Mariel Goran, a National Science Foundation IOS Award (1656510) to A.M.E. and a National Science Foundation Award (1660908) to B.M.P.. A UVM College of Medicine Shared Instrumentation Award to D.T. supported purchase of the Leica-Aperio VERSA whole slide imaging system.

## Supplemental information

**Supplemental Figure 1.**
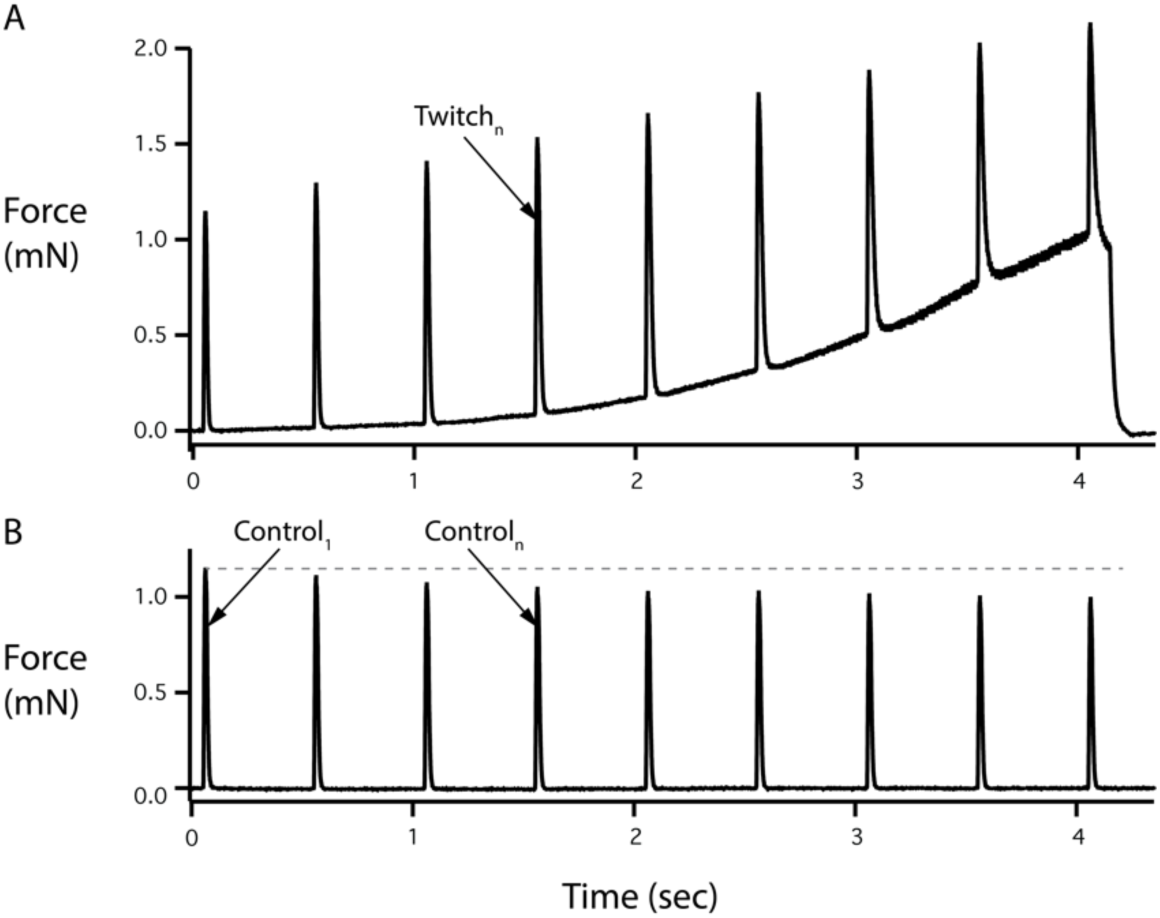
Correcting for fatigue during length: force measurements. *(A)* Representative force trace showing twitch contractions during a slow stretch from 1.00 to 1.17 x preparation starting length *(as in Main Text Figure 5B). (B)* To account for any decline in force generation associated with repetitive stimulation during the length:force relationship protocol, after a 60 second rest period, tails were subjected to the same stimulation protocol without the accompanying stretch. Active force for each twitch *(Twitch*_*n*_) in Main Text, Figure 5B, was adjusted in Main Text Figure 5C by multiplying its value by the ratio of the first control twitch *(Control*_*1*_) to the corresponding control twitch *(Control*_*n*_).

**Supplemental Movie 1.** Tetanic contraction at fixed preparation length (28°C). 1000 frames/second movie of the contraction shown in Main Text Figure 3G and 3H (50 millisecond, 300 Hz tetanus) slowed to 1/100^th^ speed. White spot in upper right corner denotes stimulation period.

**Supplemental Movie 2.** Tetanic contraction with compensatory stretch (28°C). 1000 frames/second movie of a preparation being stretched during tetanus to reduce myotome shortening, as shown in Figure 6C and 6D, slowed to 1/100^th^ speed. White spot in upper right corner denotes stimulation period.

